# Conceptual Confusion: The case of Epigenetics

**DOI:** 10.1101/053009

**Authors:** Angela Oliveira Pisco, Aymeric Fouquier d’Hérouël, Sui Huang

## Abstract

The observations of phenotypic plasticity have stimulated the revival of ‘epigenetics’. Over the past 70 years the term has come in many colors and flavors, depending on the biological discipline and time period. The meanings span from Waddington’s “epigenotype” and “epigenetic landscape” to the molecular biologists’ “epigenetic marks” embodied by DNA methylation and histone modifications. Here we seek to quell the ambiguity of the name. First we place “epigenetics” in the various historical contexts. Then, by presenting the formal concepts of dynamical systems theory we show that the “epigenetic landscape” is more than a metaphor: it has specific mathematical foundations. The latter explains how gene regulatory networks produce multiple attractor states, the self-stabilizing patterns of gene activation across the genome that account for “epigenetic memory”. This network dynamics approach replaces the reductionist correspondence of molecular epigenetic modifications with concept of the epigenetic landscape, by providing a concrete and crisp correspondence.

## 1. INTRODUCTION

Confusion arises when disparate concepts are mingled to explain or understand novel observations or when new insights undermine established concepts. This is further augmented by the ambiguity of language when distinct concepts share the same label. Ambiguity around the very term ‘gene’ and the evolution of the underlying concept is a perfect example for such confusion. Originating in the abstract description of inheritable traits when introduced by Wilhelm Johannsen in 1909, the concept ‘gene’ culminated in molecular definitions delineating transcribed and translated regions of DNA from untranscribed regulatory sequences. The molecular embodiment of a ‘gene’ as a piece of DNA, delimited in the details, however, soon ended the confusion. Awareness of the presence of different concepts independent of how we call them and of the entailed semantic and practical contradictions bears a safe discourse that allows some concepts to be unified, while others are discarded. But what if, in the absence of such awareness, different concepts are unconsciously mingled under the same term? Such is the case with ‘epigenetics’.

It is now convenient to take refuge in the province of ‘epigenetics’ – however vaguely defined – when one is confronted with the challenge of reducing the observed variability of phenotypes to a genetic explanation using the new powerful genomics technologies: the absence of “explanatory mutations” in cancer genomes, the paucity of strong genetic markers for complex diseases and the “missing heritability” of quantitative traits, in which detectable genotype variants often fail by a wide margin to “explain” the observed phenotype variation not due to environment^1-4^. But perhaps the most prosaic difficulty in mapping a phenotype to a genotype in a linear fashion is the diversity, stability and plasticity of the thousands of distinct cell phenotypes within one organism that carries one genome^1,5-7^.

The term ‘epigenetics’ has in the past decade seen a surge in popularity; mirroring the ascendance of the awareness that organismal complexity cannot be reduced to genetics (**Fig. 1**). But as diverse as the paths leading out of “genetic reductionism” are^5-10^, so is the usage of terminology in articulating alternative principles of biological information-encoding beyond that embodied by the genomic DNA. After the decades-long reign of molecular genetics that has created a tacit culture of thought in which phenotypes are by default explained by a singular, discrete material causation, typically represented by a genetic pathway, we now see the renaissance of a broader, holistic view of phenotype control, most lucidly manifest in the re-use of the visually appealing picture of Waddington’s epigenetic landscape, as in^8-14^, or the appearance of modern cartoonish variants of it, as in^11-15^. While the need for an integrative view that respects the daunting task of unraveling the intertwined molecular pathways has already been subtly articulated by the early protagonists behind the rise to dominance of the DNA as explanatory principles^5,15-17^, in reality the modern practioners of molecular biology have barely paid heed to such words of humbleness^5,16-18^.

**Figure 1.**
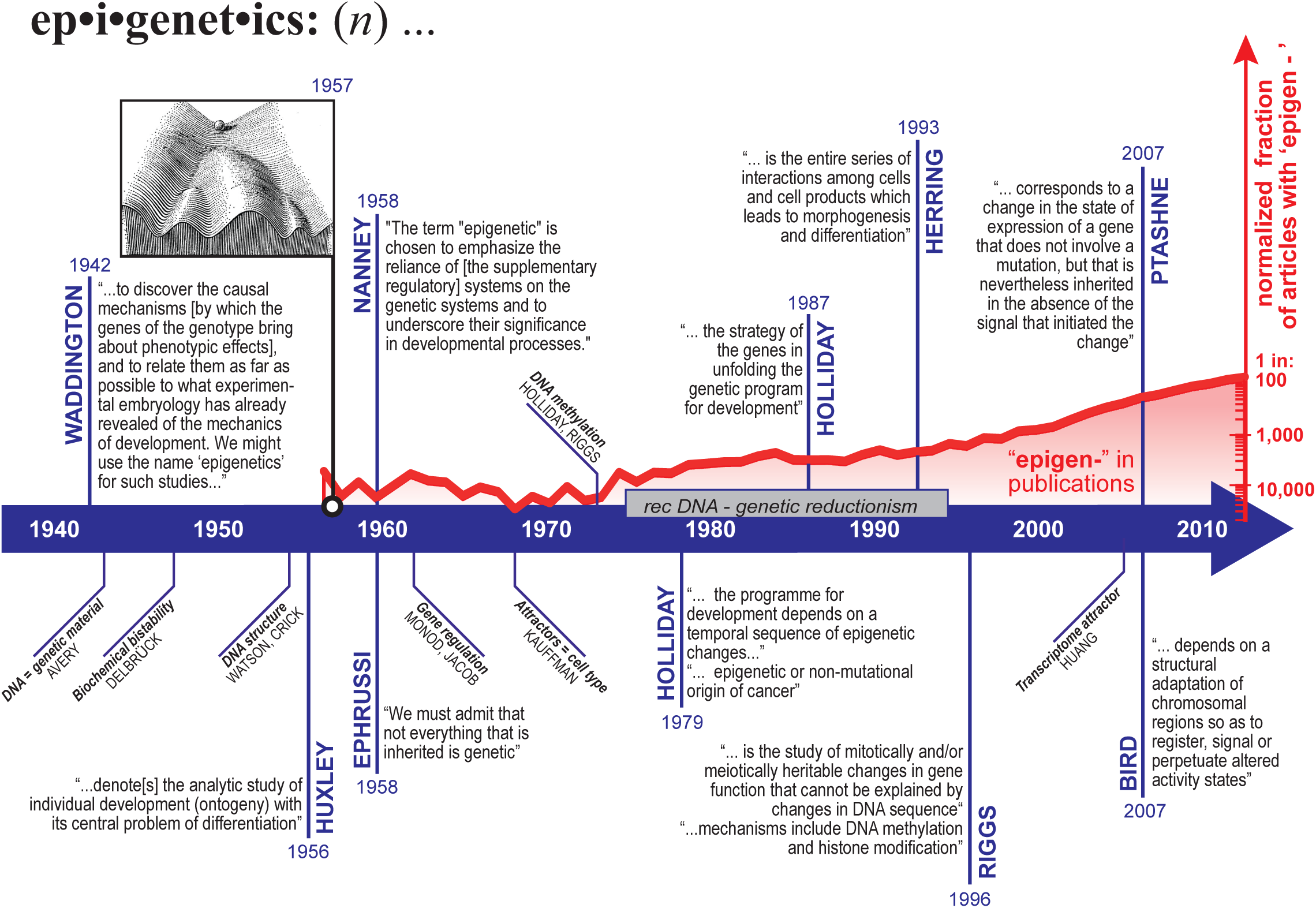
History of the term ‘epigenetics’. The discovery of DNA methylation launched the associated modern (molecular) notion of ‘epigenetics (1975) and is reflected in the surge of the number of publications after that. Note the stagnation in the two decades after that DNA recombinant technology and genetic determinism dominated mind-set in biology (grey horizontal bar). We are now seeing an enormous increase in the use of the term, as the post-genome era findings challenge the textbook gene-centered mindset. Textboxes display quotes, of selected authors (vertical text). Red curve indicate the number of Medline entries containing the wildcarded search term ‘epigen*’ per year, normalized for number of publications that contain the generic keywords ‘cell’ and ‘function’. Diagonal textboxes highlight key discoveries or findings in biology influencing the history of epigenetics but not directly related to it. The original context of the citations can be found, by chronological order, in refs. 36,37,39,50,59,60,62-64,103. See sections 2 and 3 in text for more historical details.

### 1.1. The difference between meaning and naming

So what does **‘epigenetics’** mean? And why is the concept of an epigenetic landscape so attractive? The necessity of a precise and unambiguous (however broad) definition of terms and an accurate explanation of underlying concepts for the effective communication of ideas in science is obvious. In the case of epigenetics, the desire for definitions has led to many broad ones, typically by articulating what epigenetics is *not*, namely, information encoding that does *not* involve changes in DNA sequence. But such attempts to be comprehensive are futile if one is not aware that the problem of the MEANING of a term (“What do you *mean* by ‘epigenetics’?”) has its complement in the issue of the NAMING of an entity (“How should we *name* this new phenomenon – can we call it ‘epigenetics’?”). In particular, in scientific endeavors where new phenomena are observed, novel principles discovered and innovative concepts proposed, the issue of finding a term to name any such newly described entity must receive due attention. The branch of semantics that deals with the problem of *naming* is called *onomasiology* as opposed to the more familiar *semasiology*, which is concerned with the *meaning* (“definition”) of terms^18,19^. The problem that accompanies scientific progress specifically is thus of onomasiological nature and has its starting point in a new observation or idea and not in a preexisting term. Often, new entities are first communicated by using a neologism, such as ‘transcriptome’. Alternatively, new entities are named by adopting a preexisting term. The term ‘epigenetics’ is viewed as a neologism coined by Conrad Waddington in the early 1940s, and is often interpreted as a portmanteau from the Greek preposition ‘epi’ (eni, “on” or “over”, generally indicating a superimposition) and ‘genetics’^19-21^. On an alternative account, “epigenetics” was not a completely new creation; it was derived at Waddington’s time from the already exising term “epigenesis”, used to separate developmental mechanics from preformationism. ‘Epigenetics’ thus may be seen as a contraction form reflecting Waddington’s attempt to integrate ‘genetics’ with embryology^20-22^.

The use of a preexisting term for new phenomena, if inappropriately done, has a greater potential for confusion because of the baggage of older meanings that the term carries, raising the important question: Is the new phenomenon conceptually related to the one for which the term has originally been proposed, and if so, how?

### 1.2. Two disparate major usages of ‘epigenetics’

This question becomes very concrete in the case of ‘epigenetics’ when molecular biologists, more than thirty years after Waddington created it, began to use this term to describe their discoveries. Do DNA methylation, covalent histone modifications and their downstream effects on chromatin constellation warrant to be referred by the term ‘epigenetics’ (or ‘epigenetic modifications’) simply because these modifications are on top (“epi-”) of the DNA sequence of genes (“-genetic”)? We would like to argue that the answer is ‘yes’ - in a fundamentally different and broader sense. In this modern usage of ‘epigenetics’ one rarely sees the explicit reference to Waddington’s idea where ‘epi’ is used.

In a nutshell, there are two major but disparate uses of the term ‘epigenetics’ (**Fig. 2**, major dichotomy on the left):

**Figure 2.**
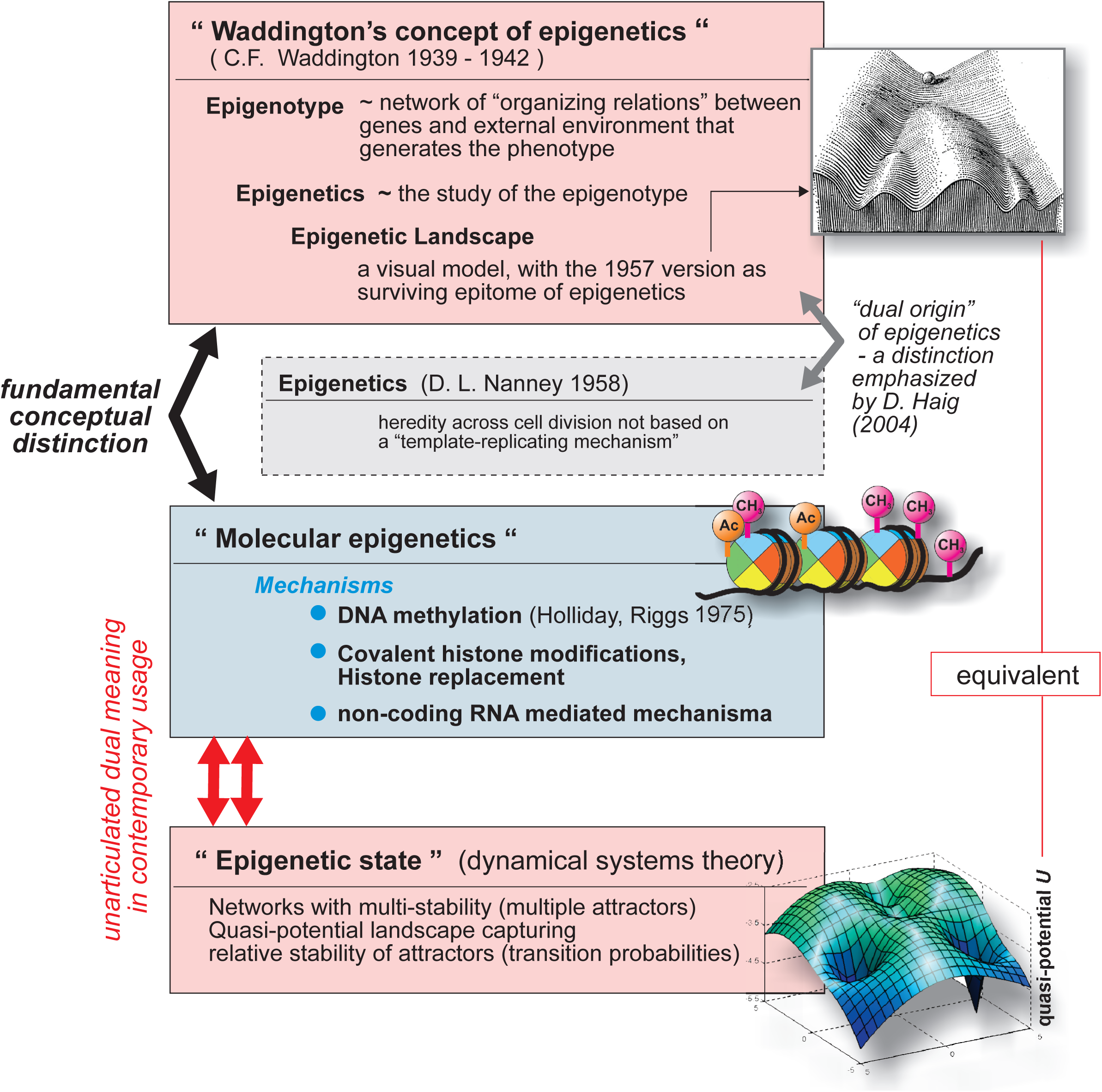
Taxonomy and dichotomies. The usage of the term ‘epigenetics’ over the past half century can be divided in two major, natural groups: usage in “*Waddington*’s sense” (top red box) or to describe a “*molecular*” entity (central blue box). These two meanings reflect a fundamental conceptual distinction: the second category is more a list of particular molecular mechanisms operating in the cell that influences gene expression state of genomic loci, while Waddington’s functional concept is broader, giving us an intellectual structure to organize and expand our biological knowledge. Separately, modern dynamical theory produced the principles of multistability in network dynamics, which applied to gene regulatory networks, leads to the notion of epigenetic state/memory (bottom red box), used mostly by systems biologists. We argue here that this third notion of epigenetics is equivalent to Waddington’s idea of epigenetics. These two concepts are epitomized in either the quasi-potential landscape or the epigenetic landscape, respectively. Haig (2004) pointed to a minor dualism in the term epigenetics. Most confusion in current communication, however, stem from the semasiological sloppiness of not separating between the material “molecular epigenetics” and the epigenetics that arise from the dynamics of regulatory interactions.

i. the more general use refers to the totality of processes “beyond” the genes that gives rise to the phenotype and goes back to Waddington and
ii. the more recent use focuses on the molecular modifications of DNA and chromatin that are thought to mediate information encoding in the absence of DNA-sequence alterations.

Although this dichotomy has not often been explicitly articulated, many sensible investigators will have encountered the confusion that it has created in daily practice. Thus it is a natural dualism of modern usage and not identical to the one discussed by Haig, which can historically be traced back to two independent explicit proposals for how use the term^8,13,22^ (**Fig. 2**, minor dichotomy on the right). Additional use of the term epigenetics in systems biology, closer to Waddington’s idea, will be discussed later (**Fig. 2**, bottom).

For practical purposes within this discourse (but not as a general suggestion) let us call the former usage proposed by Waddington (i) **conceptual epigenetics** and the latter (ii) **molecular epigenetics**, because of its dependence on a molecular substrate (**Fig. 2**). These two modes of usage have distinct histories, reflect distinct epistemic philosophies and thus, effectively *mean* different things despite the same *name*. This has led to the conflation of two fundamentally distinct concepts, even among scholars of epigenetics^8,13,23^. The use of the same term to communicate two different things, notably by two different communities, not only prevents differentiated analysis but also hampers effective communication because of unarticulated ambiguity. Unfortunately, the case of ‘epigenetics’ is a teaching example of this phenomenon.

Yet at a profound level, if we understand that Waddington’s epigenetic landscape has a formal foundation in the physics of so-called non-equilibrium systems, and actually, unbeknownst to most biologists, represents a quasi-potential function *U*(***S***) of every cell state ***S***, as we will explore below (Sections 6 & 7), then we can begin to see at the formal level the link between the two chief meanings. But this equivalence between Waddington’s conceptual epigenetics and the modern molecular epigenetics has to be articulated carefully, using principles of gene regulatory dynamics, and cannot be assumed *a priori* (as is generally done).

Thus, with this dualism as the organizing framework, and without further inspecting the etymology of “epigenetics”, the objective of this article is three-fold: (1) to provide a cursory account of the history of the term ‘epigenetic’ in biology; (2) to show that Waddington’s epigenetic landscape is more than a metaphor because it has a molecular and mathematical foundation; and (3) to convey a new taxonomy of ‘epigenetics’ (**Fig. 2**) that naturally emerges from the first principles of dynamical systems theory and from molecular biology, and that cannot be imposed by *ad hoc* definitions.

We start with the (coarse-grained) history (**Fig. 1**), which can roughly be divided into the pre-DNA and the post-DNA era usage of ‘epigenetics’. Over the past 70 years the term ‘epigenetics’ has been used in a diverse array of contexts by biologists, with diverse perspectives on living systems and with varying meaning that reflected the change of their perspective over time (**Fig. 2**). The motivation to use the term ranges from foresight to hindsight: from advocacy for embracing what in biology is beyond “genetics” (or DNA) to admittance that after all, not everything is directly encoded in DNA. The usage of the term ‘epigenetics’ has steadily increased in this time period, as plotted in Fig. 1 (number article in the Medline database per year, normalized to the estimated total number of publications in biology). Of note is the surge after the discovery of DNA methylation in 1975 (see section 3) and the stagnation during the “boom years” of recombinant DNA technology, when the enthusiasms about DNA as the code of life in the 1970s and 1980s left little room for non-genetic explanations.

### 1.3. Understanding the dichotomy: the practical importance

We would like to emphasize with this piece that awareness of the unarticulated dichotomy between conceptual and molecular epigenetics would benefit the individual researcher’s design of experiments as well as the interpretation of their results. But most importantly, words are vehicles for communicating ideas, and when ideas behinds words are not shared by the discussion partners *and* the ambiguity per se not noticed, the discussions can become meaningless without anyone realizing it because of such “meta-ignorance”, i.e. not knowing what one does not know^23,24^. Specifically, when hypothesizing epigenetic effects to describe an experimental observation, it is commonly assumed by default that molecular epigenetics is the explanation. The term epigenetics is thus employed here following the spirit in current wet labs that values the narrow understanding of proximate molecular causes But with the arrival of systems biology, an increasing number of researchers, many not biologists by trade, let alone trained in the climate of the rapid ascendance of molecular biology of the 1970s to the 1990s, will by contrast primarily regard indirect, often non-linear, effects that emerge from the complex interconnectedness of biochemical reaction networks as the foundation of the observed behavior. Such explanatory principles typically require mathematical tools to comprehend, some of which will be outlined in Section 5. In the days of interdisciplinary systems biology experimentalists will have to communicate their interpretations of ‘epigenetics’ to theorists – a dialogue dependent on sharing the same meaning behind the same name. Whether one is intellectually more at home in the domain of the narrow molecular or the broader conceptual perspective, failure to appreciate the existence of the dichotomy and consequently, the lack of curiosity for the “other viewpoint” not only will hamper effective communication but severely limit the expansion of one’s personal perspective – in an era when the molecular, dissecting approaches and the holistic, integrative approaches are so close to being meaningfully united. Toward this end, awareness and embrace of pluralism of ideas are the necessary first steps^20,24^.

## 2. WADDINGTON’s CONCEPT OF EPIGENETICS

### 2.1. Waddington’s epigenotype and epigenetic landscape

To understand Conrad Hal Waddington’s (1905-1975) motivation in creating the term ‘epigenetics’ to describe his observations and convey his thoughts at the interface between embryology and genetics, one needs to recall that at his time neither was the molecular basis of a gene known, nor was developmental biology an established discipline^20^. To communicate his idea he had to employ a new term – he was facing an onomasiological, not a semasiological situation, and addressed it with a neologism.

Waddington struggled with the attempt to unite the disparate fields of embryology, evolution and genetics. While the former was concerned with the qualitative morphological understanding of the developing embryo (epigenesis), the latter two treated the organism as a black box and, hence, took the problem of the genotype-phenotype mapping for granted. Waddington focused on development in his quest for a rational, mechanistic basis of the genotype-phenotype correspondence and in this context he first coined the notion of the ‘epigenotype’^4,19,25,26^ as a mediator between the two. This concept set the stage for ‘epigenetics’, epitomized in the landscape metaphor that Waddington utilized to explain the inherent capacity of cells to differentiate into “discrete types”^19,27^. Essentially, he equated epigenetics with development^28^, reconciling the epigenesis of the embryo with inheritance^21^. Epigenetics was, according to Waddington, to be understood as “the entirety of the developmental processes through which the genes produce the phenotype”^25^. It may be worth noting that Waddington’s approach, unlike the modern day “anti-reductionists” who readily embrace the landscape metaphor as an icon of a holistic, non-molecular view, did not appear to reflect a dismay with molecular reductionism – if such existed at the time of his early career.

### 2.2. Epigenetics, genetics and phenotype diversification

Unlike many of the embryologists of his time, Waddington sought to link (Mendelian) genes to what was known as “developmental mechanics”. Through his work in the fruit fly he was among the first to see the connection between developmental anomalies and genetic mutations^20^. Yet, it is perhaps the repeated encounters with observations that defy the Neo-Darwinian evolution of new phenotypes^21^ – incompatible with the linear one-to-one mapping from gene to trait^29-31^ – that led to the following insight: Many genes act jointly to guide development and the phenotype arises through a “network of processes” (see below) taking place amongst genes and between genes and environment^19,26^. Perhaps the most salient of such observations is the branching diversification and differentiation of a zygote into the various cell types, all of which, according to common knowledge then and now, essentially harbor the same set of genes (with a few noted exceptions). To capture this phenomenon of one-to-many mapping Waddington used the tree-like diagrams, in the context of development firstly introduced by Weismann in 1893^20^, and proposed the train track switches analogy that later evolved into the picture of the landscape of diverging valleys.

### 2.3. The early articulation of a ‘complex network’

Waddington’s notion of a “concatenation of processes linked together in a network”^19^, at a time when the equivalence between gene and DNA was not known, was a remarkable foresight given that the very idea of a *complex network* of gene regulatory actions has only found broad acceptance as a hindsight of genomic analysis. Even after Monod and Jacob showed that at the molecular level genes physically interact via their encoded products to regulate each other’s transcription and proposed small gene regulatory circuits^32^, and even after Kauffmann presented a formal treatment of complex “genetic network” in the late 1960s^33^, the concepts of networks of genes did not spread among molecular biologists. It disappeared from the radar screen of molecular biologists that directed their attention to establishing linear chains of causation to which they quickly had become habituated in the wake of the Central Dogma^31^. It was not until the arrival of massively parallel ‘omics’ technologies decades later that gene networks reappeared on the new arena paved by systems biology in the 1990s^34^. The link between *gene regulatory network* (or GRN^35^, the modern term for the older ‘genetic network’) and the epigenetic landscape is at the core of this discussion.

### 2.4. ‘Epigenetics’ after Waddington – the notion of “state”

Waddington’s notion of ‘epigenetics’ was further generalized in 1957 by Huxley as the study of developmental processes leading to differentiation^36^ (**Fig. 1**). Moving from the physiology of development to cell biology, and after the DNA in the nucleus was identified as the carrier of genetic information, Nanney proposed, at about the same time as Huxley but with scant reference to Waddington, the term ‘epigenetics’ to describe the system of extranuclear cellular heredity that was not based on the DNA sequence^37^. According to Nanney, such epigenetic control systems are auxiliary mechanisms that help to determine the specific gene expression profile of each particular cell – thus sharing the same motivation as Waddington. Importantly, as also emphasized by Haig^22^, Nanney, who criticized the gene-centered “template-replicating” concept of molecular genetics, also proposed to consider the concept of a “Steady State” by which he “*envision[ed] a dynamic self-perpetuating organization of a variety of molecular species which owes its specific properties not to the characteristics of any one kind of molecule, but to the functional interrelationships of these molecular species*”^38^.

In 1958, Ephrussi, who like Waddington had championed the idea that genetic mutations played a critical regulatory role in development, suggested that epigenetic mechanisms involve *functional states* of the nucleus and explicitly articulated the idea of non-genetic inheritance: “*We must admit that not everything that is inherited is genetic*”^39^. Such a quote must be seen in the context of the rise of molecular biology and the “Central Dogma” that helped propagate the simple concept of one gene = one protein (= one trait) and gave birth to a modern form of genetic reductionism^31^. In the ensuing climate of gene-centric thinking, a lone outlier was Sir Cyril Hinshelwood who after his Nobel award in Chemistry in 1956 devoted his scientific efforts to the study of (transient) non-genetic inheritance in bacteria. While not explicitly using the term epigenetics, Hinshelwood proposed reaction patterns, similar to Nanney’s steady states, established by intracellular chemical reaction networks within a cell that “exhibit slow reversion”, i.e., persist over multiple generations and hence can engender a memory that mediates non-genetic inheritance^40,41^. Hinshelwood’s work has almost completely been overlooked by historians of biology, let alone by its practitioners^42^; however, research along similar lines of reasoning has in the past years gained traction with the rediscovery that non-genetic persisters play a role in accelerating the evolution of bacterial resistance to antibiotics^43,44^ and of cancer cell resistance to chemotherapy^45^.

## 3. MOLECULAR EPIGENETICS

It was only in the 1970s, when methylation of DNA at position 5 of the base cytosine (5mC) in the CpG dinucleotide motif had attracted interest in the molecular analysis of cells that the term ‘epigenetics’ gained its second meaning, referring to covalent DNA modifications and, later, to histone modifications. This led to a first surge of the term in the literature (**Fig. 1**). (This trend is expected to increase in the modern day of next-generation sequencing because single DNA physical sequencing is expected to reveal new covalent base modifications of polynucleotides^46^)

### 3.1. DNA methylation and the reuse of the term ‘epigenetics’

The idea proposed independently in 1975 by A. Riggs and R. Holliday that the patterns of DNA methylation in cells are stably inherited across cell generations^47,48^ and that this could permanently affect gene expression, was too alluring to ignore: with these two properties, DNA methylation could serve as the molecular substrate for Waddington’s epigenotype and therefore, explain the conundrum of the diversification and stability of cell types that had motivated Waddington to introduce the term ‘epigenetics’. The general notion that DNA methylation silences genes^49^ still holds, although the full picture turned out to be more complex.

Holliday started around 1979 with the use of the term ‘epigenetics’ to describe this new regulatory facility afforded by the system of enzymes that regulate DNA methylation^50^. But since DNA methylation was for a period of time the sole molecular mechanism that could account for Waddington’s idea of the epigenotype, usage of the term ‘epigenetics’ developed its own dynamics and through the sheer mass of published material ‘DNA methylation’ became almost synonymous with ‘epigenetic modification’. Here is where the issues of the *name of a new observation* on the one hand and the *meaning of an old term* on the other hand, must be distinguished. Only then can one appreciate the extent of confusion caused by the borrowing of an existing term to describe a new phenomenon. It requires a speculative stretch and the neglecting of nuances of conceptualization to lump DNA methylation at one genome locus together with the old ideas of orchestrated action of multiple genes, networked processes and cellular states (as discussed later) that are visualized by the epigenetic landscape. But in defense of the modern adoption of an old term, *who* could, in view of the unique properties of DNA methylation that so obviously suggest the potential capacity for non-genetic and permanent encoding of instructions during development, resist the temptation of borrowing Waddington’s neologism?

### 3.2. Early controversies surrounding DNA methylation

In fact, molecular epigenetics, in its early days equated to DNA methylation, found fertile ground in the epistemic environment of molecular reductionism of the early 1980s because of its mechanistic plausibility and the tangible molecular substrate. Nevertheless, interest in DNA methylation waned for a while because its role in gene regulation met inconsistencies^51^. As W. Gilbert noted^52^, DNA methylation may not be essential for multicellular life because the classical model organisms (*D. melanogaster* and *C. elegans*) lack wide-spread DNA methylation in the adult^53^ and lower (but not much less multicellular) vertebrates exhibit only a fraction of DNA methylation in their genome compared to mammals. Moreover, inconsistencies as to whether DNA methylation is really suppressive in gene regulation^51^ and the failure to find mechanisms that would permit that “*alterations of methylation are freely imposable*” – the prerequisite for its role in establishing gene expression patterns conjectured by W. Gilbert^54^ – questioned the significance of DNA methylation as an epigenetic mechanism in the development of the phenotype. W. Gilbert’s requirement is an issue of eminent importance that we will visit later.

### 3.3. From DNA methylation to covalent histone modifications

The re-ascendance of molecular epigenetics was in part stimulated by the intensifying work on covalent modifications of histone proteins rather than the DNA itself, notably the methylation of lysines of histone 3 (H3) and the acetylation of arginines of H4, which impart the suppressed or the active state to gene loci, respectively^55,56^. These modifications are far more complex than DNA methylation and opened a rich field of research for exploring their structure, regulation and biological significance. The term ‘epigenetics’, originally used to describe DNA methylation, was extended to include the histone modifications. DNA methylation and covalent modifications of nucleosomes are now commonly referred to as ‘epigenetic marks’. Contemporary biologists now use ‘epigenetics’ to imply the covalent molecular marks on DNA and histones by default. This seismic semantic shift changes the intended meaning of ‘epigenetic’ from that originally proposed by Waddington^57^ and others in the 1950s and 60s (**Fig. 1**). Equivalently, in the era of omics, the term ‘epigenomics’ describes the genome-wide analysis of epigenetic marks using high-throughput technologies, such as ChIP-Seq and mass-spec^58^.

### 3.4. Modern definitions of ‘epigenetics’: extended definition (again)

The modern usage of the term ‘epigenetics’ has changed over time even within the community of investigators focusing on the molecular modifications. Holliday tried to capture the spirit behind Waddington’s quest for a general concept with his modern definition of ‘epigenetics’. The similarity between the wording employed by Waddington and later by Holliday in his more encompassing definition of 1987 is evident: “*epigenetics is concerned with the strategy of the genes in unfolding the genetic program for development*”^59^. Functional morphologists went further in generalization by defining epigenetics as “*the entire series of interactions among cells and cell products which leads to morphogenesis and differentiation”*^60^. Negative definitions that include some clause to the effect of ‘not related or depending on DNA sequence’ abounded in the 1990s and 2000s, as if competing for the most exhaustive definition. Perhaps the most current, and with respect to its modern usage most applicable, is according to Bird^61^, the definition given by Russo & Riggs in 1996^62^, defining epigenetics as “*the study of mitotically and/or meiotically heritable changes in gene function that cannot be explained by changes in DNA sequence [including] DNA methylation and histone modification*.”

The unease with the narrowness of defining ‘epigenetics’ by referring to a molecular entity other than DNA continues to shine through in Bird’s recent (2007) definition. There, epigenetics is inevitably seen through the lenses of molecular biology but the definition is augmented by a comment (which, notably, is longer than the definition itself) to ensure broad validity^63^:

> “…The following could be a unifying definition of epigenetic events: **the structural adaptation of chromosomal regions so as to register, signal or perpetuate altered activity states**. This definition is inclusive of chromosomal marks, because transient modifications associated with both DNA repair or cell-cycle phases and stable changes maintained across multiple cell generations qualify. It focuses on chromosomes and genes, implicitly excluding potential three-dimensional architectural templating of membrane systems and prions, except when these impinge on chromosome function. Also included is the exciting possibility that epigenetic processes are buffers of genetic variation, pending an epigenetic (or mutational) change of state that leads an identical combination of genes to produce a different developmental outcome. “

The attempts at transcending the molecular level view and reaching into the realm of the integrative and organismal is evident in Bird’s addendum through the reference to “state” and “combination of genes” and even “developmental outcome” which embraces the ideas behind the definitions given by Waddington or Nanney. Similarly, M. Ptashne expresses in a cogent commentary his criticism on the narrowness of the contemporary notion of ‘epigenetics’ as referring to histone modifications. He, too, recalls the notion of an enduring ‘state’: “… a *change in the state of expression of a gene that does not involve a mutation, but that is nevertheless inherited in the absence of the signal that initiated the change”*^64^. This statement captures the fundamental principle of memory shown in **Fig. 3**. Ptashne also correctly emphasizes the role of self-sustained feedback loops in epigenetic inheritance – as did Jablonka and Lamb^65^ (see below, 3.6.) – albeit in a qualitative manner and as one of several possible mechanisms. However, the dynamics of feedback regulation is a central topic to which we will return. An entirely positive definition has on the other hand been given by Bonasio, Tu and Reinberg, who define epigenetics as “*the inheritance of variation above and beyond changes in the DNA sequence*” – another extension that regards the molecular substrate as the natural starting point of considerations^66^.

**Figure 3.**
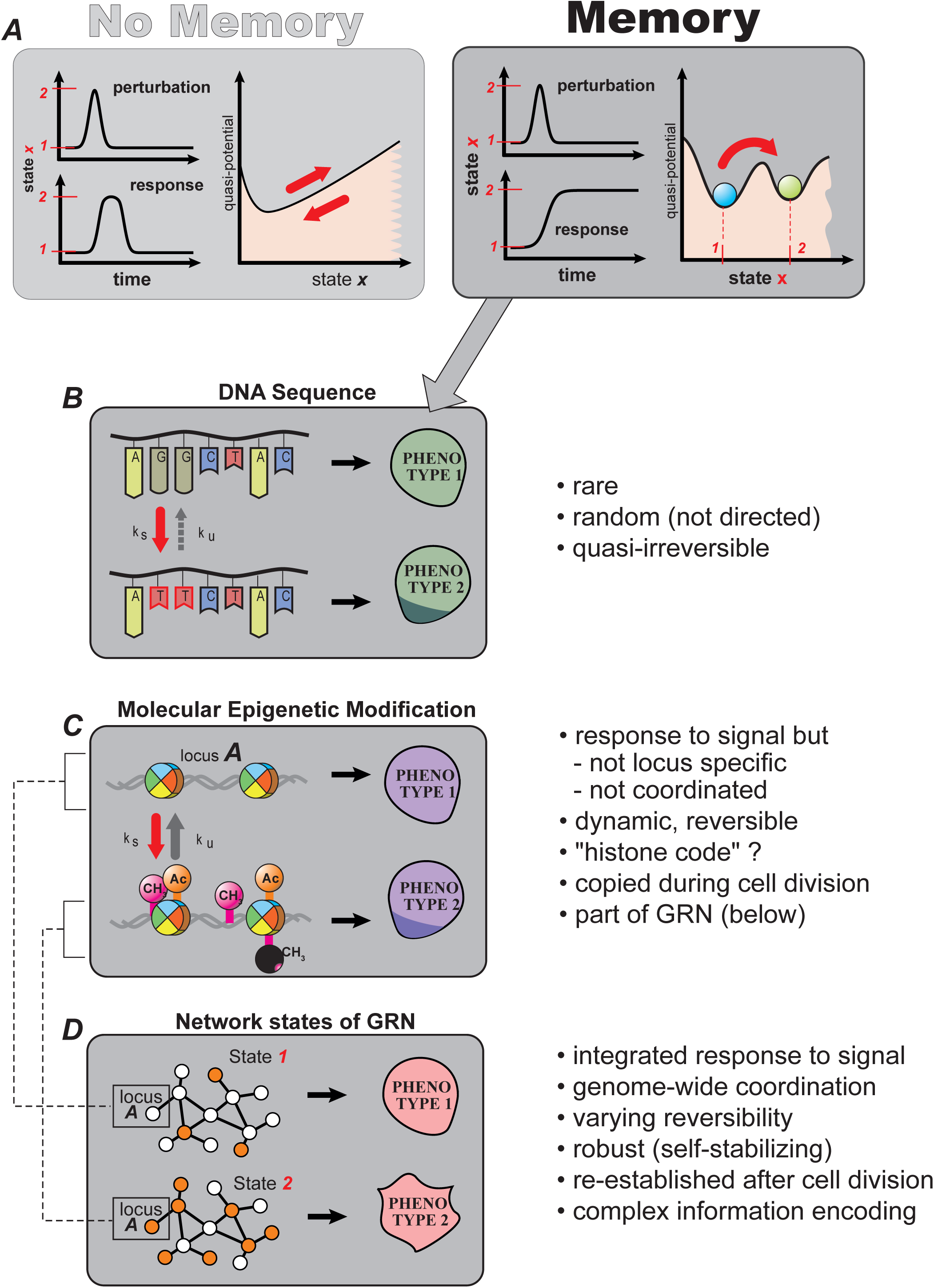
Memory in biology. Basic principle of memory (A) and various systems in biology that can encode and retain information (B, C, D). The vertical axis in (A) corresponds to a potential function (see section 7.). The prototypic memory processes include: DNA sequence changes (B), molecular ‘epigenetic’ modifications (C) and stable network states (D). The capacity of network states of “memorizing” phenotypes arises from a synergy between DNA sequence and molecular epigenetics changes. By contributing, as an additional gate, to the local (cis) control of the activity of an individual gene locus, molecular epigenetics is an elementary component of the process through which the gene regulatory network generates the epigenetic states (shown in detail in **Fig. 4**).

### 3.5. Issues with expanding the definition – closing the circle to Waddington

In all these modern definitions of ‘epigenetics’ the influence of the idea of covalent molecular modifications of DNA or nucleosomes are recognizable, but these definitons also seek to give the term a broader functional significance in order to establish universal validity. But again, not without irony (given that reference to Waddington is rarely made explicit), in doing so these definitions approach the general, conceptual epigenetics that Waddington had in mind (the epigenotype) and that Holliday himself criticized as “too broad to be useful”^67^. What we see here is the use of a semasiological approach (the extension of a definition) to address an onomasiological problem that has been perpetuated ever since it was created by calling the covalent DNA and histone modifications ‘epigenetic’. These newer definitions are welcome, but despite the re-approaching to Waddington’s ideas, they do so without providing a precise mechanism to explain how the covalent epigenetic marks on individual gene loci generate phenomena, such as discreteness of cell types, memory of entire cell states, buffering and canalization, which in the first place inspired Waddington to propose the epigenetic landscape.

In conclusion, we emphasize that the “pedestrian” use of the word ‘epigenetics’ in the modern, post-genomic era, unlike in the heydays of excitement surrounding the Holliday-Riggs hypothesis, has almost completely lost the etymological connection to Waddington’s term, let alone the connotation of his integrative concept of epigenetic. The modern (informal) usage of ‘epigenetics’ is a product of the mechanistic mindset of molecular biology and exclusively implies the molecular modifications as a phenomenon in its own right, far from suggesting the breadth of biological significance proposed by Bird, Holliday, or even, Waddington. Thus, the prefix ‘epi-’ is understood in the narrow sense of the Greek preposition for ‘above, on top’ (of DNA)^25^ – a far cry from Waddington’s notion of the ‘epigenotype’ as an encompassing category. Yet reference to Waddington is on the rise in the ‘epigenetics’ literature, even the picture of the epigenetic landscape is celebrating a renaissance - but again, in an *ad hoc*, loose and superficial manner devoid of a formal or at least precisely argued justification^8-10,12,13^.

### 3.6. Other modern uses of ‘epigenetics’

For completeness sake, we would like to mention other modern usages of the term ‘epigenetics’, which are not limited to the covalent modification, or at least place ‘epigenetics’ in a more encompassing context (**Fig. 2**). First, biophysicists have intuitively used the term epigenetics (**Fig. 2**, bottom box), hardly referring to the molecular modifications of chromatin, to imply self-sustained states in a multi-stable system (as explained below in detail). In this usage, epigenetics, as appearing in expressions such as ‘epigenetic memory’ or ‘epigenetic state’^68,69^ are functional (dynamic) phenomena and in spirit close to the principles that Waddington seemed to have intended to imply all along^70^.

A broad, colorful and spirited use of the ideas of epigenetics has been spearheaded by Jablonka and Lamb, who emphasize the multiple levels of integration in biology and evolution^71^. Although now aware of the principle by which an epigenetic state emerges from the dynamics of interactions^65^, the term ‘epigenetics’ has been employed by these authors in the sense of ‘molecular epigenetics’ to comply with the modern literature. But they have placed ‘epigenetics’ in an all compassing framework of genetic and non-genetic inheritance as a general mechanism of evolution that takes place at several levels: DNA, (molecular) epigenetics, behavior (brain, learning, culture) and symbolism (language, communication)^72^.

Finally ‘epigenetics’ is also used in conjunction with trans-generational epigenetic effects/inheritance of environmental influences, such as famine in earlier generations or in utero environmental exposure to environmental stressors^71,73,74^. The study of this phenomenon pertains to the discipline of genetics, and hence is carried out with an emphasis on the scheme of inheritance as the starting point rather than on molecular mechanisms of development. Although ‘epigenetics’ in this context should be understood in the broader meaning of Waddington^75^, DNA methylation is typically implied as the chief mechanism^76^. Here too, confusion of terminology and concepts stifle effective exchange of ideas – but this is beyond the scope of this review^71,75^.

In this broader category we may also add what selected authors consider ‘epigenetics’ in processes that to the traditional biologist self-evidently pertains to phenomena that are obviously non-genetic in nature, summarized in Table 1.

## 4. WHY MOLECULAR EPIGENETICS IS NOT EXPLANATORY

There is a tendency to invoke the molecular “epigenetic marks” whenever questions concerning how the genotype produces the phenotype cannot be answered by the demonstration of a linear causative genetic pathway.

It is conveniently assumed that epigenetic modifications of DNA and histones somehow serve as information-coding systems that account for the stability of the particular cell lineage defining gene expression patterns^77^. However, this line of thought is rooted in two old misconceptions: (i) epigenetic modifications are quasi-irreversible and (ii) they can somehow act in a gene-locus specific manner to regulate gene expression, as detailed below.

### 4.1. Reversibility of epigenetic modifications

First, the tacit notion of irreversibility is flawed because we have meanwhile learned that DNA methylation and covalent histone modifications are highly dynamic and reversible^78,79^. Despite their covalent nature, they cannot engender persistent cellular traits by themselves as required if they are to explain cellular differentiation in the sense envisioned in the 1970s, when DNA methylation first captured attention in molecular biology. The action of methyltransferases is countered by that of demethylases and histone-acetyl transferases are opposed by histone deacetylyases, etc.^80-84^ ensuring a bidirectional dynamics of epigenetic modifications. The principle *sine qua non* for a writable memory is the quasi-irreversible change of state, i.e., one that persists after removal of the agent that caused the state change and for which the reverse reaction is much more unlikely (**Fig. 3A**). Thus, the irreversibility necessary to maintain gene expression states must come from somewhere else: the gene regulatory network with its non-linear feedback loop dynamics, as we will discuss later.

### 4.2. Lack of locus specificity and the need of trans-actions for coordination

Second, the modifying enzymes that add or remove the covalent marks appear to lack locus-specificity in mammals and thus cannot produce a particular gene expression pattern that manifests the coordinated expression of a set of genes. Coordination in turn cannot be controlled by an epigenetic modification that acts in *cis* alone but requires the interaction between gene loci, an action in *trans*. Such interaction is realized because these modifying enzymes are recruited by transcription factor (TF) complexes that bind to their specific cognate regulatory regions of the genomic DNA^85^. Thus, a TF encoded by a locus *i* binds to another locus *j*, and thereby mediates the interaction between these gene loci *i* and *j*. But one may then ask: how can TFs exert prime control if their access to chromatin itself is controlled by epigenetic modification at a given locus? This chicken-egg problem is alleviated by the report that (at least for some well-studied promoter regions) the TF can bind to the target sequence even if it is in closed chromatin region and minimally exposed^86^. This is in line with the early finding that DNA methylation correlates mostly with suppression of binding of universal but not of tissue-specific TFs^49^. Perhaps the binding of tissue specific TFs to closed chromatin triggers a local self-perpetuating process that recruits chromatin modifying enzymes to fully open the locus which in turn “primes” it for the assembly of the entire transcription machinery^87^. Conversely, TF binding to heterochromatin can play a role in keeping the chromatin closed^88^. In any case, it then seems that TFs may in fact be the prime initiators of gene expression pattern: they *impose*, to use W. Gilbert’s requirement (see section 3), the epigenetic state onto a locus^54^. In fact, experiments with integrated transgenes suggest that DNA methylation is a consequence rather than cause of gene silencing^89^. Thus, as Gilbert predicted in 1985, methylation of DNA is “[not] *one of the primary top-level controls of gene expression*”^52^. In this sense, DNA methylation (and, by logical extension, histone modifications alike) should rather be viewed as “*the very bottom level, the last level of control that is used to shut genes off*” as Gilbert put it^52^.

### 4.3. Possible role of covalent epigenetic modification as an additional control layer

So what is the precise function of DNA and histone modifications if they are subordinate to the control by the transcriptional network? The fact that DNA methylation does not perform a qualitative, universal and absolutely necessary function in metazoa is consistent with its virtual absence in some multicellular species. However, it is clear that epigenetic marks exert vital functions in the quantitative tuning of expression because abrogating DNA methylation results in mid-gestation lethality^90^ and pharmacological perturbation of the balance of the modifying enzymes has substantial consequences on gene expression kinetics^91,92^ and cell fate control^93-95^. In particular, chromatin or DNA modifications appear to form local control circuits around the gene locus^96^ and to regulate the local kinetics of gene expression, e.g., by influencing the frequency of the fluctuation of chromatin between the open and the closed state^91^. These modifications influence each other at a given gene locus^97^ and may establish a localized network as additional control layer that is both subjected to regulation by TFs and, in reverse, gates their transcriptional activity.

In summary, epigenetic modifications of DNA and histones, via the modulation of chromatin conformation, play an important biological role in gene expression control far more complex than suggested by the simple notion of an array of on-off switches that constitute an epigenetic code. By themselves, epigenetic marks cannot account for the coordinated dynamics of gene expression programs that govern development.

## 5. A UNIFYING CONCEPT: EPIGENETIC MEMORY

As we have discussed above, the molecular biologist’s definition of ‘epigenetics’ can be traced back to the onomasiological error of borrowing the old term ‘epigenetic’ to describe DNA modifications. Although this term was coined by Waddington before the era of molecular genetics, it was an intuitive, rather vague association of DNA methylation with Waddington’s ideas that motivated this move. Then, using a semasiological correction to broaden the narrow equating of ‘epigenetics’ with a localized molecular alteration, the meaning of ‘epigenetics’ was expanded^63,67^ to also imply the developmental processes through which information in the genome unfolds to generate the phenotype, but remained unable to offer specific principles (Section 3).

However, a reunion of the modern notions of epigenetics with Waddington’s original ideas cannot rest on the *ad hoc* expansion of the narrow definition. One should not be misled into seeing unity when diverse groups share the same term because the same name often hides fundamentally disparate cultures of thought. Nevertheless recent developments in systems biology have paved the road to reintroducing Waddington’s 1940s proposal of epigenetics by providing a new mathematical and molecular underpinning.

### 5.1. New insights will come from dynamical systems theory

Since the early days of Waddington’s work on epigenetics, we have not only understood what the physical substrate of genesis, but we have also gained insights in the fundamental principles of the operations of complex gene regulatory networks (GRN)^35^. The theory is concerned with understanding the collective (“emergent”) dynamics in complex systems composed of many elements interacting with each other according to particular rules, of which the GRN is a prime example. Although the lessons from this young field of study have not yet attracted much attention by practitioners of molecular epigenetics, such systems can impart memory through the particular arrangement of the interactions, and hence, address many open questions of cellular regulation. Thus, applying elementary principles provided by dynamical systems theory^98,99^ to GRNs, offers new tools for analyzing the joint action of genes, as implied by Waddington, that allow us to explain in a rigorous and formal way how inheritance can be achieved without being encoded by a DNA sequence.

### 5.2. The central importance of epigenetic memory

‘Inheritance’ can be translated into a general non-biological term, suited for theoretical consideration: *memory* of a phenotype over generation boundaries (**Fig.3**). Only by embracing the principles of nonlinear dynamics can we understand in a rigorous manner how a “network of interactions”, as Waddington suspected more than six decades ago, can create memory. With such theories we will also see the connection between Waddington’s integrative concepts of “buffering” and “canalization” which cannot be accounted for by mere molecular epigenetic modifications.

Waddington’s epigenetic landscape picture was motivated by the puzzling question of how the very same genome (as we would say today) produces a diversity of cell phenotypes that are stable enough to be inherited across cell generations. We can now more concretely postulate that for such a regulated diversification of one genome into a multiplicity of cell phenotypes without mutations, a cell must harbor two capacities: (*i*) it has to remember its current phenotypic state (stability when facing noise) and (*ii*) it has to remember past encounters with (transient) external signals that changed its state (flexibility and stability of new state). Such a robust recording mechanism, or “writable memory”, is critical for producing the lasting phenotypic changes that drive development. Epigenetics, when stripped of all the grandiose definitions, is thus essentially about non-DNA based memory of a phenotypic state.

### 5.3. General ingredients of memory

For a system to have such capacity of memory, we can cast the two above requirements in more general terms (**Fig. 3A**): (*i*) it must respond to a perturbation of no relevance by ignoring it and return to its current state, which it is able to remember after being pushed out of it and (*ii*) it must also be able to respond to a signal of relevance by switching to another state and staying there even after that signal has vanished, i.e. not reverting back to the original state it held prior to encountering the signal. This plausible functional requirement for a writable memory in turn requires a set of more elementary dynamical properties of the system. It must be:

a. able to exhibit ***at least two or more quasi-discrete internal stable states*** that can ignore nonspecific influence (“random perturbation” or noise) and
b. capable of a ***quasi-irreversible state transition*** between them in response to a certain perturbation (“specific perturbation” or signal) from the outside.

Here we use the prefix ‘quasi’ because “absolute” discreteness as well as “absolute” irreversibility are not strictly required. A grey-zone is allowed, but it is much less prevalent than the black and white – or as Waddington put it: “intermediates are rare”. To illustrate this, let us consider first the case of classical “genetic memory” – the DNA and its nucleotide sequence as information coding system (**Fig. 3B**). Each sequence position can take the value of the four bases A, T, G, and C – this corresponds to 4 discrete states, between which the DNA can switch via mutations. For the sake of simplicity, we only consider substituting mutations here. A mutation of a base, e.g. from C to T, at a given position, is essentially irreversible (because the likelihood to encounter within a reasonable period of time a mutation that reverses it is vanishingly small). The conditions of ‘multiple states’ and ‘irreversibility’ are evidently satisfied. However, a switch between the states A, T, G, C occurs essentially only as a consequence of a mutation which cannot direct the change towards a particular base. Due to the non-existence of directed mutations, DNA can “remember” only random events and cannot be used to record the nature of past occurrences of a particular signal. Thus, while the genetic memory of DNA suffices for evolution following the Darwinian scheme of random mutation and selection, it is not suited for generating the controlled diversification of phenotype during development in which deterministic regulation by extracellular signals plays a major role (although selection of the fittest cells also may be involved^100-102^).

### 5.4. Enters epigenetic memory

While Waddington did not prominently address the issue of epigenetic memory, his conceptual framework is in line with our elementary notion of memory: Waddington’s concepts of “constancy of wildtype” and “buffering” corresponds to the stability of states that we suggest above, feature (a), whereas his idea of “non-genetic inheritance of acquired traits” and their “canalization” during development may be viewed as representing the writing of new information into memory and the switch to an alternative state^103^, mapping into feature (b) above.

Of course, the molecular form of epigenetics owes its ascendancy to the perceived capacity of an epigenetic mark to act as a memory device: a chromatin modification could be written by a signal that “somehow” activates the modifying enzyme to exert a change in DNA methylation status at a genomic position or in one of the histone modifications and thereby affect gene expression (**Fig. 3C**). These covalent modifications were thought to be inherently stable; however, the compelling idea of Riggs’ “secondary code”^52^ or the more recent “histone code”^104^ fall apart as explained above because these modifications neither pass the test of irreversibility nor can one fathom a specific mechanism by which any signal *Y* that is to silence or open a locus *X* through epigenetic modification can do so selectively. In addition, chromatin regulation by covalent modifications itself represents a complex local subnetwork involving crosstalk between various levels of modifications, as well as the control of this local network by trans-acting regulators^105^.

We discussed above the molecular and logical rationale for the requirement of a network of trans-acting regulatory activities that coordinates the loci in order to establish enduring gene activation patterns. But how does the gene regulatory network produce the capacity to encode input signals and memory (as indicated in **Fig. 3D**)?

The principles through which epigenetic modifications are linked to the network-dependent generation of memory are explained in a qualitative, simplified cartoon in **Fig. 4**. In the following sections we will first show how “epigenetic memory” emerges from the dynamics of the GRN (Section 6). Second, we will step beyond traditional dynamical systems theory and briefly sketch the outlines of the newer theory of quasi-potentials, which provides a formal justification and quantitative basis for Waddington’s epigenetic landscape metaphor (Section 7), schematically summarized in **Fig. 5**.

**Figure 4.**
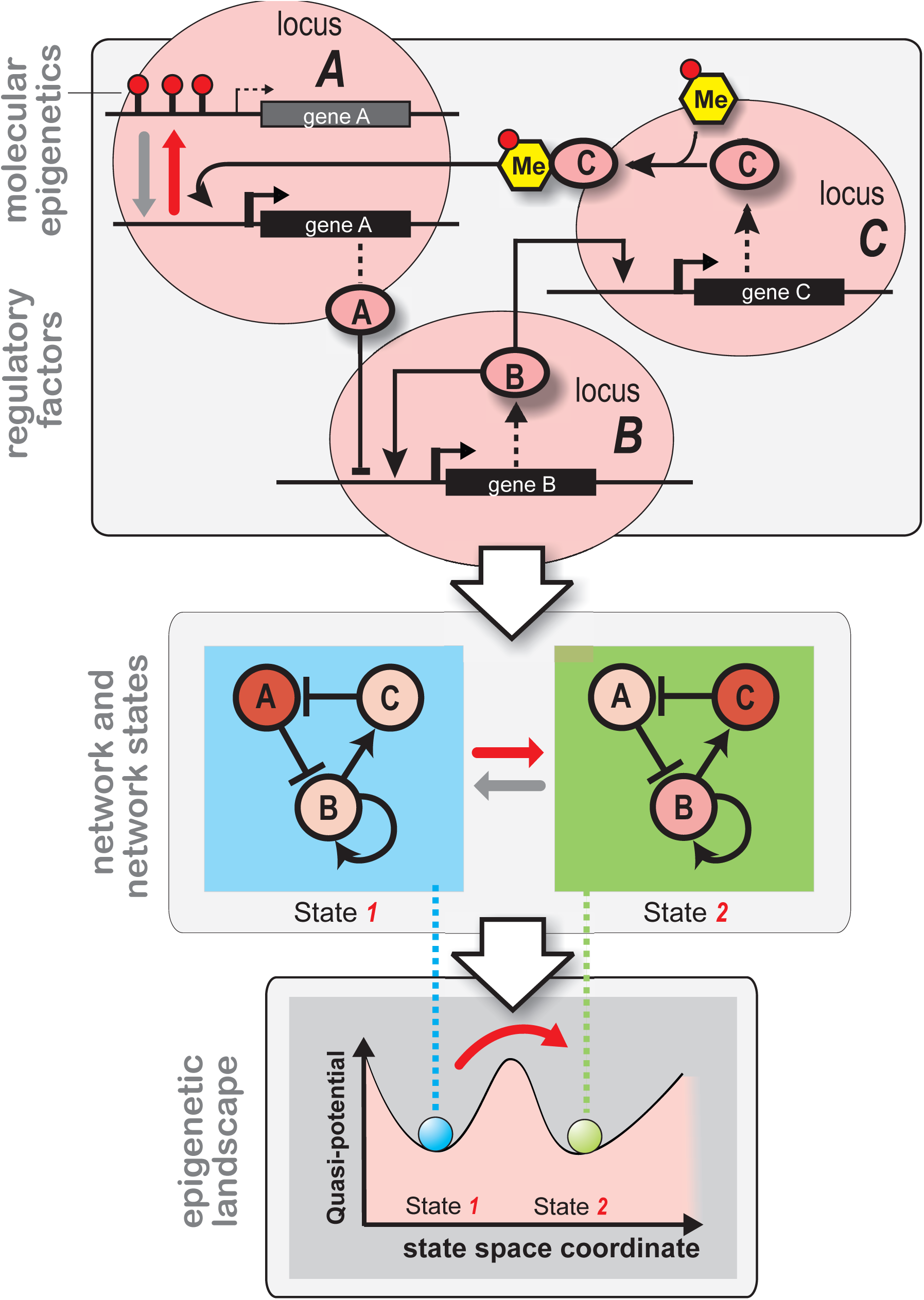
Connection between epigenetic modification and epigenetic states/landscape. The various ‘states’ of a given gene regulatory network constitute ‘epigenetic states’. They are determined by the totality of the activity of each of gene loci that participate in the regulatory network. Since epigenetic modifications affect the activity of a gene locus they contribute to the epigenetic state of the network. Herein lies the connection between ‘molecular epigenetics’ and ‘epigenetic states’. *Top panel*: A three-gene-loci (*A, B*, *C*) network. For locus *A* two different states of epigenetic modification are shown (red lollipops indicate DNA methylation). Note that these *cis*-epigenetic marks must be imposed by a factor acting in *trans* (from another or the same locus): in this case, locus *C* encodes a transcription factor that binds to the promoter region of locus *A* and recruits a DNA methyltransferase (‘Me’, yellow) there. Black arrows represent regulatory interactions, grey and red arrows in locus *A* represent biochemical reactions (transitions between conformations). Each locus (large pink ellipsoids) represents a node in the regulatory network. *Middle panel*: Coarse-grained view of the same network in which each locus is shrunk to a network node (circles *A*, *B*, *C*). This higher-level view displays the network transition between two network states, 1 and 2. The shade of the nodes indicates the activity of each locus as a result of the regulatory interactions (= network vertices, black arrows). Note that the epigenetic state is a property of the *entire* network, defined by the activity levels of each node and is indicated by the blue and the green background. Again the grey and red arrows represent transitions, now between the entire network states. *Bottom panel*: Each network state has a given “stability” (see text, section 7 and **Fig. 5** for details), such that stable states are attractors (valleys) in a quasi-potential landscape (epigenetic landscape). This establishes the final leg in connecting the molecular biologists’ epigenetic modification with Waddington’s epigenetic landscape.

**Figure 5.**
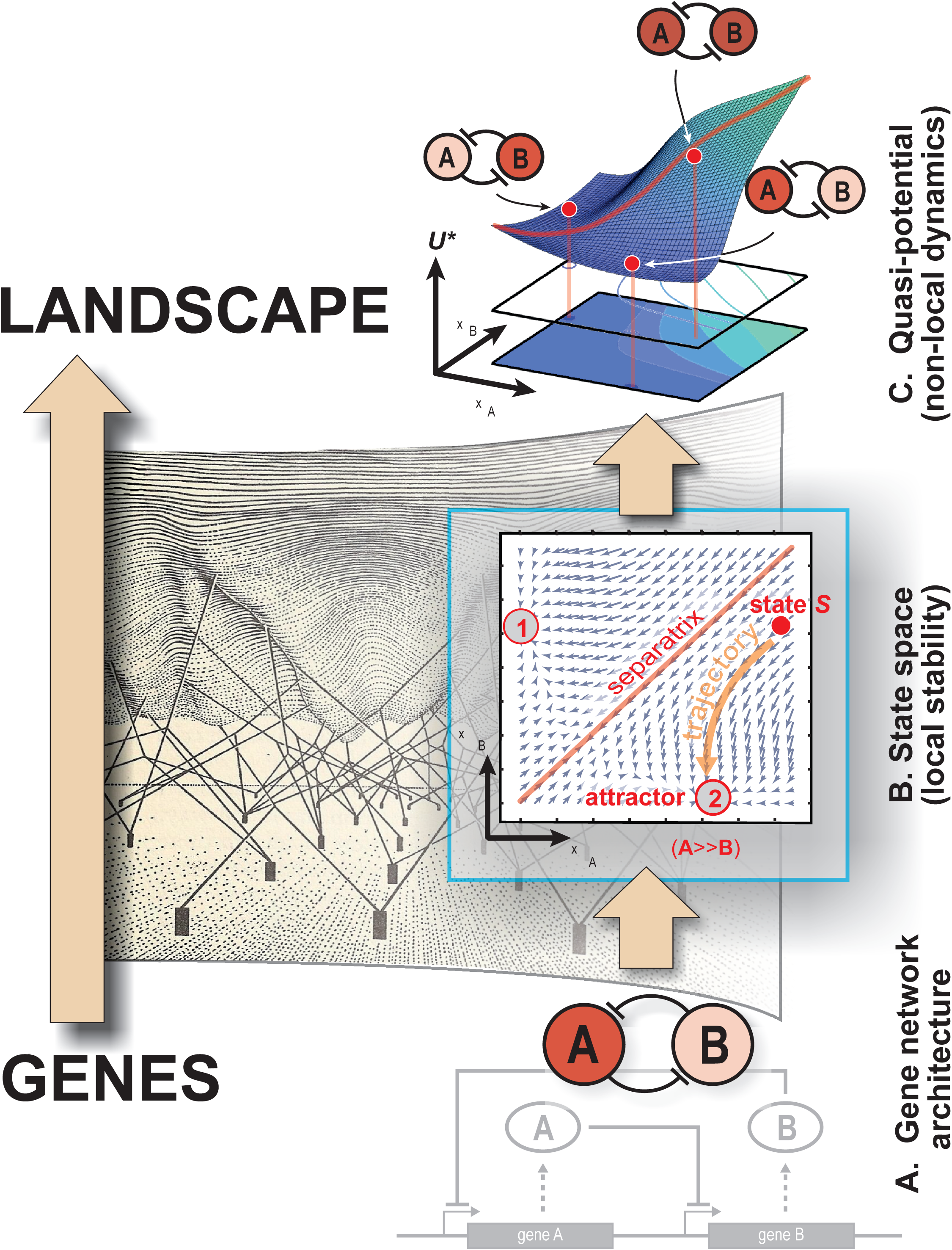
From regulatory network to quasi-potential. Waddington’s conceptualization (backdrop, left half) of a mechanism defining the shape of the epigenetic landscape put in context of the modern perspective: a web of genes (dark rectangles) anchored on a fixed plane defines the epigenetic landscapes by concerted action (joining lines), hauling down ponds and chreods (adapted from Waddington, 1957). The modern concept – illustrated for a simple two-gene system, right pane – makes a congruent and rigorous connection from the architecture of gene regulatory networks (A), via the traditional state space (B) to the description by a quasi-potential landscape (C).

## 6. MEMORY WITHOUT DIRECT MOLECULAR ENCODING: ATTRACTORS OF GENE REGULATORY NETWORKS

For the gene regulatory network to develop memory capacity it must exhibit multiple quasi-discrete states. In fact, some gene regulatory networks with a particular “wiring diagram” exhibit precisely such property, which is well known to occur in non-linear dynamical systems and is referred to as ‘**multi-stability’**. One of the simplest cases is the so called bistable switch, a mini-network consisting of two regulatory genes (typically, transcription factors (TF)) that regulate each other in a mutually repressive manner: TF *A* suppresses expression of TF *B* and vice versa (**Fig. 5A**). Such mutual repression circuits occur naturally and are often encountered in key cell fate decision circuits (although embedded within larger gene regulatory networks).

### 6.1. The dynamics of the bistable two-gene circuit

Let us focus for pedagogical purposes on the two-gene circuit. Because the two genes of this mini-network inhibit each other’s protein expression, the system typically exhibits two stable states (assuming that the magnitude of the strength of repression, basal activity and natural inactivation are within some range). In state 1, the TF *A* is very highly expressed and *B* is lowly expressed, in brief (*A*>>*B*), while state 2 has the opposite “gene expression pattern” (*B*>>*A*). Even without equations the integrated network behavior can be plausibly explained in a qualitative manner: The more highly expressed one of the two genes (e.g., *A*) is, the more does its protein repress its own repressor (in this case *B*) which in turn further increases its own expression (*A*).

This positive feedback loop, composed of a series of two inhibitions, drives a cycle propelling the increase in the expression of gene *A* (and decrease of B) until further increase of *A* is balanced by the natural inactivation of *A*, the rate of which is typically proportional to the actual expression level (“first order decay”). At this point increase and decrease of A are balanced (at a high level) and the system enters in a stable stationary state (steady state) defined by the expression pattern (*A* ≫ *B*). By symmetry the same principles of course can be applied to show that the configuration (*B*>>*A*) is also stable. Because the driving force towards these two stable, stationary states emanates from a self-perpetuating positive feedback loop and vanishes only when the circuit is in one of these two steady states, they are robust to minor “disturbances” by external factors that may affect the expression of these genes.

By contrast, the balanced state (*A*=*B*) is a stationary state but it is unstable because any deviation of it will trigger the positive feedback loop, pushing the system towards either one of the two stable states, (*A*>>*B*) or (*B*>>*A*) – depending on the ‘direction’ of push of the disturbance. The two stable steady-states are called “**attractor states**” because any configuration of the system (which in reality we can measure as the expression pattern of *A* and *B*) that is not exactly (*A*=*B*) will be attracted to either state, (*A*>>*B*) or (*B*>>*A*) where the circuit reaches a steady-state. The co-existence of these two stable stationary states in a particular system earns it the attribute “bistable” – a specific case of multistability. So far, qualitative concepts to explain such self-maintaining stability of distinct states have centered around the notion of “self-sustaining feedback loops”, an intuitive principle which in the era of molecular epigenetics only few have articulated^64,65,71^ and which Waddington was increasingly aware of, but did not explicitly describe in his earlier work.

### 6.2. A minimal formalism to describe the dynamics: states and their space

The above dynamics can be more formally studied by the mathematical description of this system as a non-linear dynamical system for instance in the form of *ordinary differential equations* (ODEs). We introduced above the state of a (2-gene) network, which we call ***S*** (defined by the expression levels of the genes *A* and *B* of the network at a given time). What the ODE does is to describe the “driving force” ***F*** that acts on the state ***S*** to change it. The direction and magnitude of the force itself depends in a particular fashion on the current state ***S** (the relative levels of gene expression A and B)*, thus we write ***F***(***S***) (**Fig. 5B & C**) – for an introduction see e.g.^1,99^. How ***F*** depends on ***S*** is determined by the network and is what is described by the ODEs. In other words, any given state ***S*** – except a steady state – experiences a force ***F*** produced by the collective action of the regulatory interactions of the network. This is the reason why the notion of a ‘state’ – which is defined by the gene activity patterns across a set of genes or the genome and is measured, e.g., as the protein abundance in the cell, is of central importance. Here, the state ***S*** is defined by the levels *x*_*A*_ and *x*_*B*_ of expression of genes *A* and *B* at a given time *t*: ***S***(*t*) = (*x*_*A*_, *x*_*B*_).

The collective behavior of the genes *A* and *B* results from the constraints imposed by the network (**Fig. 5B**) of trans-effects such that their change of expression (the change of values of *x*_*A*_ and *x*_*B*_) is not independent from each other but coordinated, as dictated by the network architecture and captured by the ODEs. The constraints imposed by the network on the change of the state ***S*** are manifested in the force ***F***(***S***) = ***F***(*x*_*A*_, *x*_*B*_) that prescribes how *x*_*A*_ and *x*_*B*_ have to change given their current values or given the current state ***S***(*t*) = (*x*_*A*_, *x*_*B*_). If we now view (*x*_*A*_, *x*_*B*_) as a pair of “geographical” coordinate values, then the network state ***S*** is a point at position (*x*_*A*_, *x*_*B*_) in the ***state space*** of the network – the space spanned by the two dimensions that represent the expression levels *x*_*A*_ and *x*_*B*_ of the genes *A* and *B*, respectively (**Fig. 5B**). Each point of the state space is defined by the gene expression pattern (*x*_*A*_, *x*_*B*_) associated with the respective state ***S*** at a given time. In other words, each gene expression pattern, or network state ***S***, maps into a specific position in the state space. This mapping is at the core of visualizing the network’s dynamics: an abstract “state” is translated into a concrete “position”. Correspondingly, a change d***S*** of the gene expression state ***S*** maps into a change (d*x*_*A*_, d*x*_*B*_) in its positions, which is a movement of the state ***S*** along a given ***trajectory*** that reflects how *x*_*A*_ and *x*_*B*_ change in a particular interdependent way. This movement reflects the network constraints: the trajectory is preordained by the regulatory interactions of the network through the position-dependent force ***F***(***S***). In the bistable switch case, every starting point ***S***_0_in the state space has thus a predestined endpoint – one of the two attractors (circles with red numbers 1 and 2 in **Fig. 5B**). If one were to paint all points ***S*** that end up in the same attractor state with a given color, coherent regions of the state space of the same color will emerge that can be referred to as a ***basin of attraction*** of that respective attractor.

The state space view is a powerful visual representation widely used in dynamical system theory (**Fig. 5B**). It captures the constraints on the collective dynamics of the genes that is coordinated by the network. One can depict the forces ***F*** at selected initial points ***S***_0_ (e.g., placed in a grid) as vectors indicating where the respective state would move in the next time unit under the regulatory influences and constraints of the network, from (*x*_*A*_, *x*_*B*_) to (*x*_*A*_+d*x*_*A*_, *x*_*B*_+d*x*_*B*_). This gives rise to a ***vector field*** that carries all information on the dynamic development of a system as defined by the ODEs – hence, by the network. Its graphical representation can be regarded as a snapshot of the dynamics of a system, akin to a photo of moving objects taken at longer exposure time to visualize the traces of motions. The state space is compartmentalized by the ***separatrix***, which in the case of two attractors is a line delineating the boundaries of the basins of attraction.

### 6.3. Memory and attractor transitions

With this example of bistability, we have now satisfied the two conditions for memory: the existence of multiple states and the fact that each one is a stable attractor. These states are stable in the sense that in response to any small perturbation **Δ*S*** on ***S*** that kicks the network (or cell) out of the attractor state 1 (by changing the expression levels of genes *A* or *B*), the system will relax back to the attractor state 1 as long as the perturbation is small enough so that the perturbed state ***S***+**Δ*S*** does not cross the boundary of the basin (separatrix). Conversely, if the perturbation is sufficiently large (typically, due to a specific combination of change in both *A* and *B* expression) then the network may land on the other basin of attraction and thus be attracted to the alternative attractor state 2. Importantly, because of the intrinsic stability of attractors, once the perturbation has vanished the network will still reside in attractor 2, thus meeting the second condition for a writable memory.

Moreover, the attractor formalism also offers a straightforward explanation for non-genetic inheritance of acquired traits that was central to Waddington’s studies. If genes *A* and *B* are transcription factors in the cell which resides in a given attractor 1 then a cell division will cause a minor disturbance, a perturbation of the state from ***S*** to ***S***’, i.e. a small change of the expression values (*x*_*A*_, *x*_*B*_) due to the random, non-perfectly symmetrical partitioning of the cellular proteins to the two daughter cells. For specific network architectures, as is the case of the bistable switch, such a perturbation will not allow the cell to switch to the other attractor. In the state space depicted in **Fig. 5B**, cell division amounts to a perturbation with an ensuing movement of the current state ***S*** which for an attractor typically cannot cross the separatrix, unless the unequal partitioning exceeds a certain level. Thus, the memory of a gene expression pattern that is encoded by the attractor, which in turn is a product of the regulatory network, can be preserved across cell divisions and, in principle, no epigenetic modification of chromatin is needed.

### 6.4. Extension to more complex networks: The case of critical dynamics

Although this pedagogical example utilizes a 2-gene network, the dynamical systems concepts introduced here are valid for networks of *N* genes, leading to a *N*-dimensional state space. Inspired by the work of Waddington and the discovery of gene regulation by Monod and Jacob, Kauffman pioneered in the late 1960s the study of complex networks of hundreds of thousands of genes in computer simulations, elegantly circumventing computational limitations by using Boolean functions for interactions between simple “on-off” genes to model complex GRNs^33,106^. Kauffman demonstrated the stunning intrinsic capacity of a defined (but large) subset of such complex, even randomly wired networks, to produce, despite the hyperastronomical number of possible states and trajectories, not chaos but a countable number of stable attractor states^107^. The collapse of complexity and emergence of ordered yet non-rigid behavior, which he termed “critical dynamics”, naturally explains the intrinsic facility of genomic networks for both stability and flexibility and hence, to generate an ordered diversification of phenotypic states into a large number of distinct, stable variants that would represent the cell types.

We know now that the mammalian genome encompasses 2000-3000 or so gene loci for regulatory factors, such as transcription factors and non-coding, regulatory RNA that control the expression of a defined set of other loci, and that it generates a repertoire of perhaps thousands to tens of thousands discrete cell types or phenotypic states. Gene expression profiling technologies that measure the transcript levels across the genome now offer means to monitor the trajectories of the network state ***S*** in cells (although transcriptome measurements only represent an approximation of the activity status of the gene loci). Yet, such simple transcriptome dynamics measurements permit us to demonstrate how thousand-dimensional trajectories spontaneously converge to stationary gene expression profiles, providing the first molecular level evidence for the existence of point-attractors in the high-dimensional space of gene expression and their correspondence to physiologically distinct phenotypic cell states^108,109^

### 6.5. History of the idea of cell types as attractors

The concept that the discrete cell types of the metazoan body are attractor states of an underlying molecular network has been proposed for a long time, but this natural idea went into oblivion with the ascendance of molecular biology and its habit of linear, mono-dimensional causation. Its dominance in biology fostered an enduring climate of thought in which biological observations are almost by default reduced to the one-gene-one trait principles. DelbrÜck in 1948^11^0, Jacob and Monod in the 1960s, soon after their discovery that genes regulate genes^32^, Thom in the 1960s, based on mathematical considerations, apparently inspired by Smale to introduce the term “attractor”^111^, and Kauffman in 1969^33^ all proposed, in some form or another, the general idea that cell types are the manifestation of attractor states of a network of “trans”-regulatory molecular interactions.

Even without the mathematical and molecular framework, we would argue that Waddington’s thinking, although communicated in descriptive terms and through metaphors, can be considered entirely congruent with the formal description of state space and attractors. He was the first to recognize the rather abstract properties of stability and discreteness of cell types - which are ontological features of attractors, using phenomenological descriptions, such as “constancy of the wild-type” and the “sharply defined tissues” and “rarity of intermediates”^1927112^. He also had a robust notion of trajectories which the cell has to follow “by necessity” - because they are imposed by the regulatory constraints of the network. Waddington introduced the concept of ‘chreods’ - an apt blend from Greek for ‘necessary’ (xpsia)) and ‘path’ (656) to describe what we know as trajectories in state space. In his last (unfinished) book, “Tools for thought” (published posthumously), Waddington recasts his idea of epigenetics in terms of dynamical systems concepts and complex systems sciences along with new versions of his iconic sketches^70^.

Waddington’s awareness of the attractor idea can be traced back to the legendary “Towards a Theoretical Biology” meeting series that he organized at the Villa Serbelloni in Bellagio (Italy) in 1965-1968. Here he explicitly articulated the connection between chreods and trajectories in the basin of attraction: “*I have proposed the name ‘chreods’ for such a multidimensional domain which contains a vector field converging on to a time-extended attractor*”^28^. This insight was stimulated by his encounter with René Thom^113^ and Stuart Kauffman at these meetings. It was here that Waddington first saw in Kauffman’s ideas that attractors of random Boolean networks represent cell types and that transitions between these attractors represent pathways of cell differentiation – a possible gene network realization of his epigenetic landscape (S.K., personal communication).

While the correspondence between network attractors and the concrete image of valleys in the epigenetic landscape is appealing and intuitively plausible, a rarely articulated conceptual barrier that needs to be addressed separates these two ideas. To achieve this in a formal way entails an extension of the above dynamical systems approach into the domain of non-equilibrium and stochastic dynamics and the concept of quasi-potentials. In the following we can only touch upon these ideas briefly but hopefully with sufficient detail to just permit a qualitative comprehension of the first principles that connect specific gene regulatory network features with the specific topography of Waddington’s epigenetic landscape.

## 7. FROM STATE SPACE TO QUASI-POTENTIAL AND (BACK) TO THE EPIGENETIC LANDSCAPE

Waddington himself sensed the link between epigenetics and networks, albeit at a more intuitive level and articulated in words and pictures but not in mathematical concepts (see above section). A perhaps subliminal awareness of networks was already reflected in his early usages of expressions, such as “processes linked together in a network” or “set of organizing relations”^114^, in his attempts to bridge the conceptual gap between genotype and phenotype. His 1958 depiction of ropes tied at one end to pegs underneath the landscape and at the other end pulling the landscape at distinct points to produce the valleys^27,112^ is a first sign of the intuition that somehow the landscape emanates from the collective action of genes via a set of general laws (**Fig. 5**, backdrop).

### 7.1. The limited analogy to potential functions: the idea of quasi-potentials

To all who are familiar with classical physics and chemistry, the sight of Waddington’s landscape picture will strike the resonant chord of “energy potentials”, notably the concepts of potential wells and energy barriers between them that explain the separation of states, their stability and the effort it takes to transition between them.

There is something to this analogy; but there are crucial and profound differences that are often overlooked. Thus, a few words of caution – without entering the technical discussion of a still evolving field – should be in order. One central caveat is that gene regulatory networks are open systems far from thermodynamic equilibrium that continuously consume (dissipate) energy and require continuous inflow of energy to maintain its order^115,116^. Hence, it is formally not correct to speak of ‘potential energy’, or just ‘potentials’, which are properties of equilibrium systems. The vertical axis, or the “elevation” in Waddington’s landscape, does not represent potential energy or free energy as in mechanics or in chemistry. Thus, it is often and more appropriately referred to it as ‘*quasi*-potential’ – or a generalized potential.

### 7.2. Intuitive utility of the landscape picture: Relative stability, Global dynamics and Cell lineage switch

Why do we need a potential landscape? The state space with its vector field description and its structures, the attractors and their basins (**Fig. 5C**) capture the entire repertoire of the dynamical behavior preordained by the gene regulatory interactions. What is the added value obtained from a global landscape description? While attractors are well-defined elements of the integrated dynamics of networks, namely, as configurations representing stable steady states, they are defined *locally*, independent from each other. However, in a multi-stable system we have an interest in relating the various attractors to each other in terms of their “relative depths”^115^ and the relative heights of the barriers that separate them. This is because in developmental biology the processes at the center of interest span an entirely different scale, a much broader range of the state space, covering several thousands of attractors, than in most engineering applications of the attractor concept, where one is concerned about local deviations from one single attractor state of a system (e.g., stable flight or steady temperature). Development is a long sequence of signal-triggered or noise-induced transition between many stable and metastable attractors in which the proliferating cell “swarm out” in state-space to occupy the various attractors to assume the stable gene expression patterns of the individual cell types^117^.

Waddington’s epigenetic landscape is appealing because it subliminally offers the observer such a comparison of the various attractors. The smooth hills of varying height between the valleys stimulate the thought of jumping between valleys – corresponding to switching cell lineages – a possibility that Waddington considered in view of his developmental biology experiments^20^. The landscape imagery becomes relevant when one considers transitions between the attractors (valleys) – due to random disturbances (“noise”) or controlled perturbations.

### 7.3. Generating the landscape from the state space – the problem in the analogy

To visualize the quasi-potential landscape we return, for sake of simplicity, to the 2-dimensional system (2-gene network) whose state space is represented by the XY plane (**Fig. 5B**). Then the third, vertical dimension *Z* can be used to depict Waddington’s elevation, let’s call it now *U*_wad_, that is assigned to every state ***S***, such that we have the scalar function *U*_wad_(***S***) whose graphical representation is a landscape surface over the XY plane (**Fig. 5C**). For *N*-dimensional systems (*N*>2, in fact *N*=1000s of genes), the concept of assigning a quasi-potential value *U* to each position of the network’s state space position is still valid, but if visualization is the aim, we would have to perform a transformation to reduce dimensionality: Assume, e.g., that the entire *N*-dimensional state space is compressed (“projected”) in some (non-trivial) way into the two dimensional XY plane, such that neighboring points on this plane still preserve (as best as possible) the neighborhood relationships they had in the *N* dimensional space^118,119^.

The landscape graph offers a first visual impression of relative depths of attractors: it is harder to climb out of a “deep” valley than a shallow valley. Similarly, to use Waddington’s description, the “undulated surface” is, as a whole, “tilted” to capture the overall directionality of development^120^. Thus, one expects that the elevation Z represents some kind of global potential energy, as is familiar from classical mechanics, experienced when rolling massive objects on a rigid surface in the presence of gravity (Waddington’s marble): It is expected that they would seek the point of lowest elevation or “potential” energy *U*_wad_ (bottom of valleys) that is accessible from a given position. More importantly, one would expect, by analogy, that the elevation differences between state 1 and state 2 would immediately tell us whether we can move without net energy expenditures (i.e., spontaneously) from state 1 to state 2, no matter what energy barriers lies in between them. From here we see that the landscape suggests a behavior independent of the actual path in the state-space.

Such an interpretation of Waddington’s epigenetic landscape as energy landscape, based on the intuition from dealing with idealized, frictionless conserved systems is not correct. The concept of a potential function *U*(***S***) is useful when it can be linked to the driving forces ***F***(***S***) (see section 6, **Fig. 5C**) through the gradient of *U*(***S***), or ∇*U*(***S***). The gradient ∇*U*(***S***) can be imagined for the two-dimensional system as the direction of the steepest descent of the landscape at position ***S***. By computing the gradient one obtains the driving force at a given position ***S***, i.e., ***F***(***S***) *=* –∇*U*(***S***) in the case that the landscape topography, given by the function *U*(***S***), is known rather than the mechanisms (=network architecture) that generate the forces ***F***(***S***) at each point ***S***. This inverse procedure, starting from the landscape to determine the trajectory, is of course what the rolling marble does for us. However, it requires that we know the shape of the landscape, the function *U*(***S***).

But here is the *nervus rerum* that causes headache: for almost all higher dimensional (*N* > 1) system, there is no systematic way to find *U*. Even worse: a potential function *U* such that ***F***(***S***) = –∇*U*(***S***) does not even exist in general, i.e., ***F***(***S***) is non-integrable. The non-existence of a potential function in the classical sense means that going from state *1* to state *2* is path-dependent since energy conservation does not apply^121^. This is in principle comparable to the real world situation where the mechanistic details of the motion matter: climbing up a mountain along a steep bumpy road may take more energy than walking along a longer but smooth designated trail to the same point. This path-dependence essentially implies that one cannot actually define a global potential function that consistently relates different positions on the landscape to each other.

### 7.4. Noise-based attractor landscape

Yet, if there is no global potential function *U*(***S***) why does Waddington’s epigenetic landscape not only appeal intuitively but also appear to capture the key features of developmental dynamics so well? There has been many attempts to compute with some approximation a “quasi-potential” function *U* in view of the non-integrability of ***F***(***S***)^68,122,123^. A simple and widely used approach is to model the dynamics of a “noisy” system. Gene expression noise^124,125^ is inevitable due to the stochastic nature of transcription and protein synthesis and causes gene expression levels in each dimension to fluctuate randomly. In such systems the networks (cells) do not move along smooth trajectories but “wiggle” around, even within attractor states, such that the nominal attractor state position ***S*** only represents an “average” position. Because of the stochasticity, one can naturally assign to every state a probability *P*(***S***) of finding the system in that particular state ***S***, given some level of gene expression noise, when steady-state is reached. Such steady-state probability *P*(***S***) is inversely proportional to the stability of a state (the more stable a state is, the more likely the network is to be found there). Hence, one is tempted, inspired by Boltzman, to define a probability based potential *U*_prob_, as the negative logarithm of the respective probability, *U*_prob_(***S***) = – log *P*(***S***)^126^. This translates the relative stability manifested by the steady-state probability into the landscape elevation *U*_prob_(***S***) at point ***S***: the lower the elevation *U*(***S***), the more stable is that state. Therefore, attractor states are located safely at the bottom of potential wells. However, this notion of a potential based on noise and probabilities does not capture the truly global dynamics, since in order to probe large-scale topography structures, the noise must be high enough to jump over large hills separating distant attractors, but at the same time this can blur the local landscape structures.

### 7.5. Formal utility of the landscape picture: attractor transition probabilities

In contrast to the desired “global potential”, whose definition for a non-equilibrium system, such as the GRN, is tricky, and perhaps impossible, a quasi-potential in the sense of a Wentzell-Freidlin potential Δ*V*_12_ can be exactly defined^121^. This potential is derived from “the least action path” between two distinct attractor states **1** and **2**. Carrying on the mountain analogy, the quasi-potential is now implicitly defined through the path along which least effort is required to hike from one valley (attractor) to the other. Such information is sufficient to infer the transition probability for going from attractor **1** to **2** and thus captures a notion of the “relative depth” of the two attractors. This Wentzell-Freidlin potential Δ*V*_12_ can be linked to the driving force ***F***(***S***) of the system that we encountered above as the “force” in the vector field as follows, for details see^127^.

As previously discussed, ***F***(***S***) is in general a non-integrable (path-dependent) field, so a potential difference between two points in that field cannot be obtained by computing the “path integral” of the force between the points **1** and **2** – for any path the integration yields a different result. However, ***F***(***S***) in most general terms can be decomposed as ***F***(***S***) = –∇*U\** + ***F***_r_ with ∇***F***_r_ = 0, the sum of two-component vector fields such that the first of them is still integrable: (1) the gradient ∇*U*\* of a scalar field *U*\* and (2) the remaining vector ***F***_r_, which is *not* the gradient of a potential. Thus, ***F*** is not a gradient field as long as ***F***_r_ does not vanish. But if the decomposition is chosen such that the gradient force ∇*U\** is perpendicular to the “rest-force” ***F***^127^ then the potential difference of this function between points **1** and **2**, *ΔU*_12_*, can be shown to be (up to a constant factor) identical to Δ*V*_12_ (Freidlin and Wentzell, 1984). The scalar field *U*\* can thus be used to depict a landscape of some interpretational value: *U*\* now captures the “non-local dynamics” that encompasses the regions relevant to describe the transition between two given attractors (“notch between two valleys”). Again, however, one should not forget that the non-gradient component ***F***_*r*_ in ***F***(***S***) = –*∇U\** + ***F_r_*** in general does not vanish, i.e., ***F***(***S***) ≠ –∇*U*. We do not obtain a “global” scalar potential that would permit us to compare the elevation between multiple distant points *ad libitum*. An alternative decomposition has been presented by Ao and colleagues which is equivalent to the one presented above^128^.

Therefore, strictly speaking, a true *global* potential *U* does not exist to warrant the energy landscape analogy. However, one can define relative elevations ***A**U* that suffice for the purpose of capturing *“nonlocal”* properties of the dynamics, namely when one considers the transition between a specified pair of states. In this sense, the landscape picture conveys information that the state space description of a vector field picture does not provide.

Conversely, the *local* structure of the landscape defined by ***F**(**S***), in particular close to the boundaries of a basin of attraction, where cells fate decisions occur, reflects the dynamics of attractor transitions accurately^129^. It is thus tempting to speculate whether genomes may have evolved to encode GRNs of a very specific type of architecture which allow it to govern the development of multi-cellular organisms via the successive branching of chreods, while optimizing the balance between *stability* of cell phenotypes and *ability* to transition or “jump” between adjacent lineages when necessary.

## 8. CONCLUSION: UNIFYING MOLECULAR EPIGENETICS AND WADDINGTON’s EPIGENETICS

(1) We have started our tour of the many meanings of the term ‘epigenetics’ with Waddington’s ideas; we then discussed the diversity of concepts behind it and the change of vantage points over the past 70 years, tracking the rise of molecular reductionism – only to return to Waddington’s landscape. But we revisit Waddington’s concepts equipped with a new set of theoretical tools that allows us to recognize the formal link between the modern, molecular notions of ‘epigenetics’ and Waddington’s integrative epigenetics. In summary, the three major concepts behind the term ‘epigenetics’ are depicted in **Fig. 2**. In solving the riddle of the many meanings of ‘epigenetics’ we first linked molecular epigenetics to gene regulatory networks and its dynamics. Then, we linked the non-local dynamical behavior of a gene regulatory network, manifest in attractors and attractor transitions, to the quasi-potential landscape. The latter formalism offers a measure of direction and probability of state transitions and thus, of the relative stability between two attractors. Once presented with the mathematical derivation of the quasi-potential landscape as a graph that captures the dynamics of the molecular interactions, it is not difficult to see the correspondence of the quasi-potential landscape to Waddington’s metaphoric epigenetic landscape – thus closing the circle. We can now propose that the epigenetic landscape, or for that matter, the gene network’s quasi-potential landscape, is the central, unifying concept of the various forms of epigenetics^31^.

(2) But how useful is the landscape idea for the practicing biologist? By establishing the molecular and mathematical foundation of the landscape image, we free it from the suspicion of being a pedestrian, post-hoc, mental model molded to fit the observation, and lift it onto center stage of a formal theory that is solidly grounded on first principles of mathematics and the material reality of gene regulation networks. The equivalency of the two landscape ideas shines a new light on Waddington’s true achievement. The utility of the landscape model for biology is immense. We have only scratched its surface in this piece whose primary goal was to address the semantic confusion behind ‘epigenetics’. Waddington utilized the metaphoric landscape with much effect to illustrate a variety of concepts and to explain observations that defied Neo-Darwinism and the central dogma, such as canalization, buffering, non-genetic inheritance, genetic assimilation, phenocopies, the Baldwin effect, etc. These phenomena that so naturally and intuitively are accounted for by his landscape can now, owing to its equivalency to the quasi-potential of network dynamics, be mapped back to the framework of gene regulatory network and hence, linked to molecular processes.

(3) Concretely, we can now sharply distinguish between lasting genetic and environmental changes, yet understand their effects within the very same model: mutations in regulatory genes change the network architecture (because they ultimately result in altering the way a trans-acting gene interacts with its targets). The hence rewired network architecture, according to the network dynamics model, will have changed the topography of the landscape^130^. As a consequence, chreods may be redirected, access to attractors that are important to organismal fitness may be closed down, or previously hidden attractors may be made accessible, paving a new developmental path to a novel complex and stable variant phenotype that can now be exposed to natural selection. In contrast to mutations, small transient perturbations do not alter the genome and hence leave the network architecture unchanged. Therefore they do not alter the landscape but push the cells (Waddington’s marbles) around in the valleys, occasionally deviating their course to alternative valleys, thereby triggering a new lasting phenotype. Enduring environmentally induced but not genetically encoded changes ensue.

(4) The landscape picture of the genome opens a new vista, which was previously blocked by the monolithic neo-Darwinian ideology and the emphasis on molecular epigenetics (“epimutations”), onto the interface between genetic and non-genetic factors and between evolution and development. These dualisms, for which there has been no shortage of unification attempts, can now be studied under a single conceptual framework.

## ACKNOWLEDGEMENTS

The authors thank the Institute of Systems Biology for support and Dr. J.X. Zhou and Dr. A. Skupin for fruitful discussions. A.O.P. is funded by BBSRC, S.H. and A.F.d’H. are funded by the NIGMS Center for Systems Biology (2P50 GM076547), S.H. is Alberta Innovates the Future Scholar.

